# Normalizing need not be the norm: count-based math for analyzing single-cell data

**DOI:** 10.1101/2022.06.01.494334

**Authors:** Samuel H. Church, Jasmine L. Mah, Günter Wagner, Casey W. Dunn

**Affiliations:** Department of Ecology and Evolutionary Biology, Yale University, New Haven, CT, USA; Yale Systems Biology Institute, Yale University, New Haven, CT, USA; Department of Obstetrics, Gynecology and Reproductive Sciences, Yale Medical School, New Haven, CT, USA; Department of Obstetrics and Gynecology, Wayne State University, Detroit, MI, USA

## Abstract

Counting transcripts of mRNA is a key method of observation in modern biology. With advances in counting transcripts in single cells (single-cell RNA sequencing or scRNA-seq), these data are routinely used to identify cells by their transcriptional profile, and to identify genes with differential cellular expression. Because the total number of transcripts counted per cell can vary for technical reasons, the first step of standard scRNA-seq workflows is to normalize by sequencing depth, transforming counts into proportional abundances. The primary objective of this step is to reshape the data such that cells with similar biological proportions of transcripts end up with similar transformed measurements. But there is growing concern that normalization and other transformations result in unintended distortions that hinder both analyses and the interpretation of results. This has led to an intense focus on optimizing methods for normalization and transformation of scRNA-seq data. Here we take an alternative approach, by avoiding normalization altogether. We abandon the use of distances to compare cells, and instead use a restricted algebra, motivated by measurement theory and abstract algebra, that preserves the count nature of the data. We demonstrate that this restricted algebra is sufficient to draw meaningful and practical comparisons of gene expression through the use of the dot product and other elementary operations. This approach sidesteps many of the problems with common transformations, and has the added benefit of being simpler and more intuitive. We implement our approach in the package countland, available in python and R. By explicitly considering counts in terms of their measurement process, we avoid and overcome many challenges in modern RNA-seq and open new avenues for the analysis of these data.

## 1 Introduction

Counting transcripts in high-throughput assays is now both routine and highly productive for many lines of research^1,2^. Next generation RNA sequencing assays, including single cell RNA sequencing (scRNA-seq), measure the abundance of gene transcripts across samples^3,4^. In most assays, sequencing reads are collected, mapped to known genes, and measurements are reported as read counts per gene in a given sample^2^. When RNA samples are drawn from bulk tissue, the number of transcripts is typically large enough that counts for all but the rarest transcripts can be considered as measurements of proportional abundances and described with rational numbers^2,5^. As such, counts for each sample are often normalized to sequencing depth, scaled by a multiplier (e.g. 10,000), and log-transformed^6^. In recent years, single-cell and single-nucleus mRNA sequencing (scRNA-seq and snRNA-seq) have become practical and commonly applied assays^3,4^. Where bulk tissue RNA-seq projects typically measure RNA for a handful of samples, scRNA-seq data are generated for thousands of cells in a single experiment. However, because the total counts per cell are much smaller than the counts per sample in bulk experiments, there are severe problems with interpreting counts as measurements of proportional abundances^7^. This is because for typical scRNA-seq data, the vast majority of count values per gene and sample are zero, and the majority of non-zero counts are very small^8^.

The first step in most scRNA-seq analyses is to move data out of count-space by rescaling and transforming values. In the process, the measurements are transformed from natural numbers into rational numbers (numbers with fractional parts). The motivation of these steps is to reshape the data so that cells with similar biological proportions of transcripts end up with similar transformed measurements^6^ and fall closer to each other in the multidimensional space of gene expression. However, a growing chorus has raised warnings that these transformations lead to unintended distortions^7,8^. One of the primary issues is that stochasticity dominates the sampling process in this low-count regime, so that for many genes observed counts are a poor proxy for proportional abundance^7^. Because logarithmic transformations cannot be applied to zero values, additional steps of adding arbitrary pseudocounts (e.g. +1 to all values) are common, but this can introduce new biases^7,9^. Furthermore, rescaling low count values can invoke a false sense of confidence in relative gene abundances. From a philosophical point of view, transformations that lead to negative or fractional counts, that have no physical interpretation, violate the central tenet of measurement theory that data should correspond to the physical reality they represent^10^.

Though problems with transforming scRNA-seq data out of count-space have been well described, there is little consensus on how to remedy them. Recent suggestions include embracing zero values by converting single-cell data to binary zero/non-zero values^11^, invoking alternative transformations (e.g. square root, proportional fitting) for variance-stabilization^12^, or avoiding the use of log-transformations by modeling single-cell data with a Poisson measurement model^7,13^. Here we argue that there is great value in treating the data as what they are – counts of transcripts sampled from the total population of mRNA – rather than as estimates of proportions. We present and evaluate a new analytical approach that preserves the original measured properties of the data by considering them in a vector space over natural numbers (see Appendix 1), rather than transform them into rational numbers. We demonstrate that a restricted algebra that is closed over the natural numbers, requiring no scaling or transforms that would result in values other than natural numbers, is sufficient to perform many of the key tasks in scRNA-seq analysis. These tasks include assessing cell similarity, measuring differential gene expression, and identifying clusters of cells with shared signatures of expression.

We present our implementation of count-based scRNA-seq analysis in the `python` and `R` package countland, available at github.com/shchurch/countland. The software seeks to complement existing tools that are currently vital to scRNA analysis (e.g. `scanpy` in `python`^14^, `Seurat` in `R` ^15^), by offering an alternative approach that sidesteps common data transformations. We then test this software on benchmark data and show that this approach can accomplish the standard objectives of scRNA-seq analysis, avoid pitfalls associated with transformation-based workflows, and improve the interpretability of results.

## 2 Building intuition for count matrices

By increasing our familiarity with the structure and composition of a count matrix we can build our intuition about how common transformations are likely to reshape it. Count matrices of scRNA-seq data encode the number of counted transcripts for each gene in each cell (Fig. 1). Here we show cells as rows and genes as columns, though conventions vary (typically, cells are rows in `python` and columns in R). Because values within the matrix represent observations of physical transcripts by sequencing instruments, they are always either zero or positive integers.

**Figure 1:**
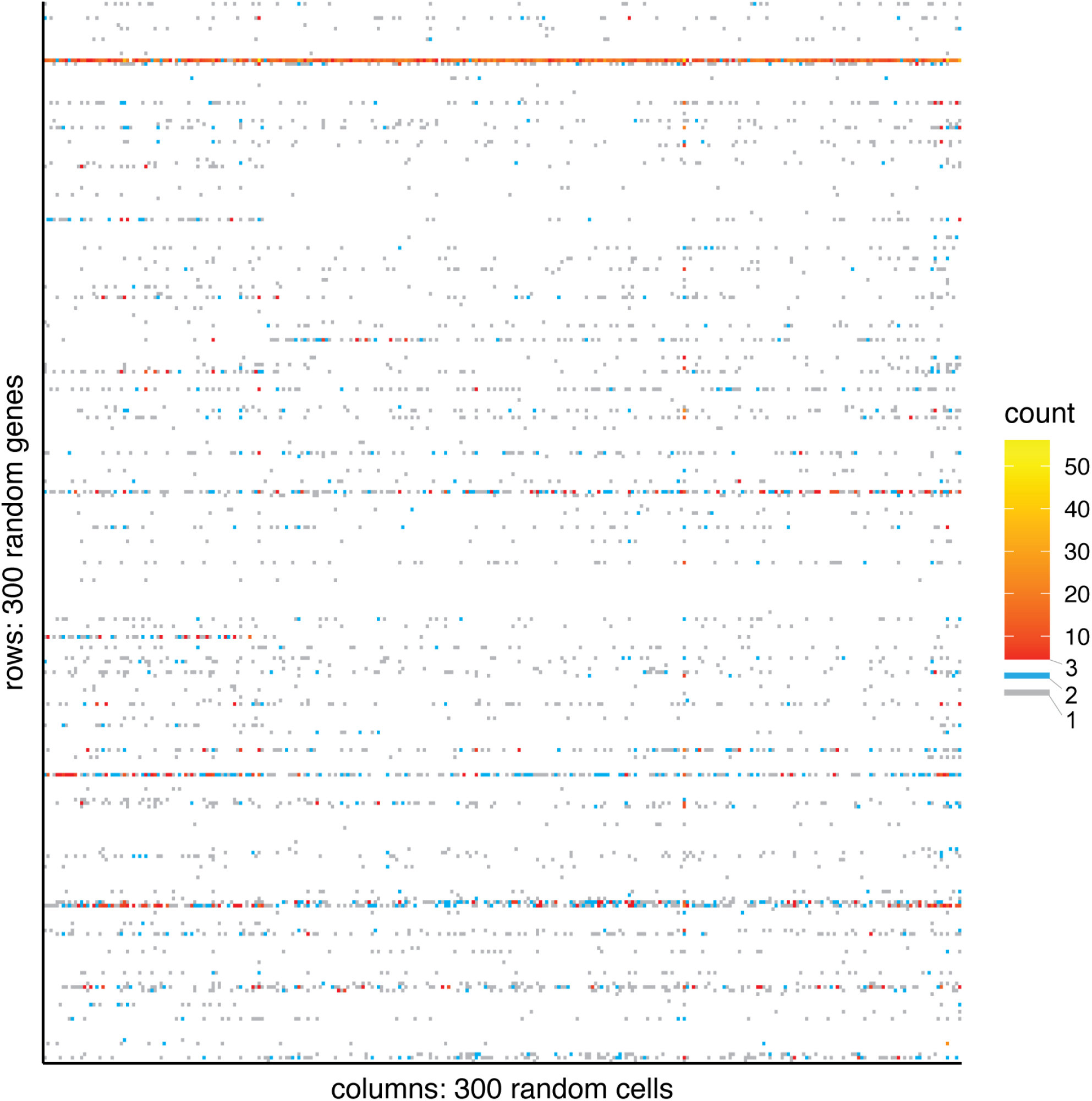
Single-cell RNA count data are mostly zeros and small integers. Visualizing a portion of the count matrix from the Silver standard PBMC dataset, described by Freytag *et al* (2018)^16^. After count values of 0 (white), the most common values are 1 (gray) and 2 (blue). Count values of 3 (red) and higher (yellow) are rare and concentrated in a few genes with relatively high expression.

Visualizing the raw count matrix shows that the dominant feature is the presence of a large number of zeros (Fig. 1). These zeros can be considered as having three possible origins: biological, sampling, or technical^17^. Biological zeros are the true absence of a given transcript in the cell, and may be the result of a gene not being expressed in a given cell or all transcripts of a gene having been degraded by the time of observation. Sampling zeros arise because we count only a small fraction of all transcripts in a given cell, so by chance many genes that have transcripts in the cell are not detected^18^. Technical zeros arise from artifacts that eliminate counts for some genes in some cells, and can be introduced during cell preparation, mapping, or other analysis steps. Several modern techniques such as using unique molecular identifiers (UMIs) have helped bring technical zeros to within distributional models of sampling^19,20^. Increased sequencing depth can reduce the number of sampling zeros^18^. Despite this, the benchmark for scRNA-seq is a data matrix with >90% of values zero, compared to 10-40% in bulk RNA-seq experiments^18^.

Of the remaining non-zero values, most are of very small magnitude. In the widely used PBMC3k benchmark dataset, for example, 69.8% of non-zero values are exactly one and 88.6% are less than five (Fig. 1). This means that for a given cell or gene, it is often true that there are no or very few values greater than two across all measurements. Cells and genes with count values greater than ten are the exception rather than the rule.

The fact that count matrices are dominated by zeros and low integer values has implications for how we attribute meaning to the data. RNA sequencing seeks to profile the proportional abundances of all transcripts at a given instant in time, but our instruments do not directly measure proportions. Instead, with scRNA-seq we sample a small fraction, usually around 10%, of the total transcripts per cell^4^. It is generally assumed that the probability of sampling a read is determined by the proportional abundance of the transcripts of each gene^18^. Normalizing the data to the total counts for each cell (the first step of a standard workflow) invokes an additional assumption that sampling has been sufficient such that observed counts are a robust proxy for proportional abundance. But at low magnitudes, sampling one additional transcript results in big jumps in the apparent relative abundances. Adding a single read that increments a count from one to two, for example, results in a two-fold increase in proportional abundance, and incrementing a count from zero to one results in an infinite-fold increase in proportional abundance.

Consider a cell that contains 10,000 transcripts, comprising 1,000 genes in different proportions. If we sampled all 10,000 transcripts, then counts would be a perfect proxy for proportional abundance, and the difference between a gene with one transcript and a gene with two would constitute a two-fold increase. But if we sampled only ten transcripts, counts would be a very poor proxy for proportional abundance. In this case, a likely scenario is that we would observe one count each for ten genes, but that does not indicate that each of those genes is present as 10% of total transcripts. Furthermore, the likelihood of a second observation of a gene would increase when the relative abundance is larger, but does not give us a precise numerical estimate for the fold difference between genes. Though this example is extreme in that only 10 transcripts are sampled, the fact that real count data contain mostly zeros, ones, and twos suggests the same principle applies to the shallow scRNA sequencing process.

In summary, the problem with rescaling count data can be stated as follows:

- The presence of a large number of zeros, ones, and other low integer values indicates a shallow sampling process in which stochasticity is a major factor.
- At low integer values, large jumps in relative abundance are common (e.g., a 100% difference between one and two), and are disproportionate to the biological reality of transcript abundance.
- As such, counts relative to sequencing depth are a poor numerical proxy for proportional abundance of transcripts.
- The practice of normalizing by rescaling count data (natural numbers) to proportions (rational numbers) bakes these stochastic sampling effects into the data by assuming that sampling is sufficient such that counts are a robust proxy for transcript proportions. From then on, counting effects cannot be deconvoluted from biological signals.

The problem with other common transformations include:

- Log-transformations stretch the differences between small integer values. Because differences between small integers are frequently influenced by the sampling process, these transformations exaggerate stochastic sampling effects over biological signals in the data.
- The dominant feature of the data is the large number of zeros. This kind of dataset is not a good candidate for log-transformation, where zeros have no numerical interpretation.
- While adding pseudocounts can make calculation practically possible, it is *ad hoc*, introduces its own biases, and has no correspondence to the physical data or underlying sampling process.

## 3 A restricted algebra for counts

### 3.1 Count-space

In mathematical terms, scRNA-seq count measurements occupy a high-dimensional vector space over the natural numbers (here defined as inclusive of zero, see Appendix 1). These are computationally encoded as unsigned integers. There is no such thing as a negative or non-integer transcript count, as these are physically meaningless.

Group theory is the field of modern algebra that concerns collections of numbers and operations on those numbers. In group theory, a group is defined as closed when specified operations on members of the group result in objects that are also members. Groups can then be classified by which operations they are closed for. One familiar type of group is a field, which is closed under the operations addition, subtraction, multiplication, and division. Rational numbers form a field because applying any of these operations to rational numbers always results in a rational number. Natural numbers, such as transcript counts, do not form a field, because natural numbers are not closed under subtraction or division. These operations can result in values that are outside the group (e.g. non-natural numbers like -2 or ½). Groups that are closed under addition and multiplication but not under subtraction or division are classified as semirings. See Appendix 1 for a formal treatment of the topic.

Constraining operations to those that are closed over a semiring provides a restricted algebra for analyzing scRNA-seq data that maintains its original count nature, guides intuitive thinking, and is still surprisingly rich. To understand what operations in the space can tell us, we can visualize observations of transcripts in cells as vectors in count-space. In count-space each gene forms an axis, perpendicular to all other gene axes. This space, therefore, has as many dimensions as there are genes. Each sequenced cell exists at a point in count-space with coordinates given by the counts for all genes. This point is the terminus of a vector emanating from the origin, which is the point where all gene counts are zero. We can conceptualize the cell sitting at the origin, for which all counts are zero, as the null cell.

The measurement process can be thought of as estimating this vector by walking through count-space from the null cell (Fig. 2). As RNA sampling proceeds, each new transcript moves the cell away from the origin, always by one integer value in the positive direction parallel to the axis of the corresponding gene (Fig. 2). The final number of steps is determined by the sampling depth for the cell in question, and is the total number of counts for a cell.

**Figure 2:**
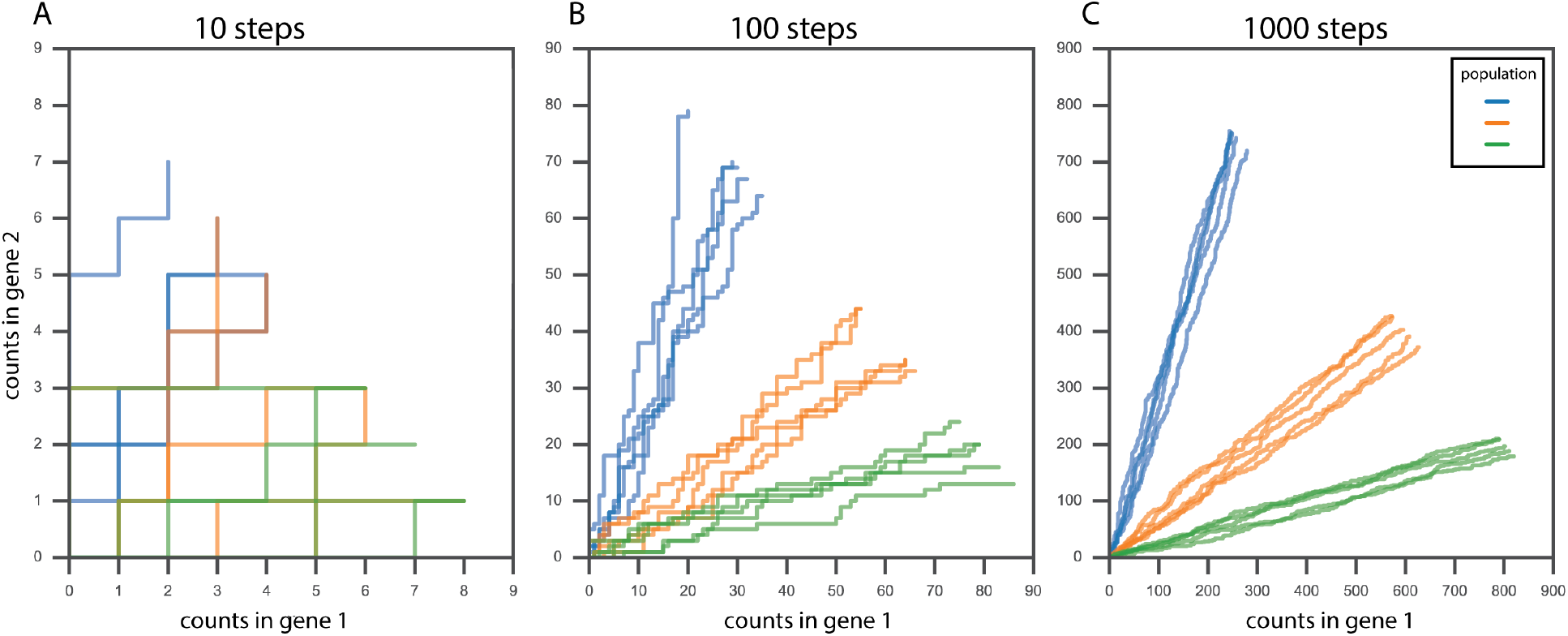
A simulated RNA counting process in a system with two genes and three cell populations with five cells each. One step equals one new observation of a transcript and moves the cell one unit along the axis of the corresponding gene. Colors indicate cell populations. A, Paths after counting ten transcripts. Note that after an equivalent number of counts, all vectors terminate along a diagonal line with a slope of -1. B, Paths after counting 100 transcripts. Vectors begin to separate based on the actual proportions of transcripts in different cell populations (different colors). C, Paths after counting 1000 transcripts. Vectors corresponding to different cell populations terminate in distinct regions of count-space, and vectors from the same population are more similar to each other than to vectors from different populations.

In this count-space, there is no concept of Euclidean distance. Calculating Euclidean distance requires the operations of subtraction and square roots, under which the natural numbers are not closed. Visualizing vectors in a reduced two-dimensional count-space helps clarify why Euclidean distance is not the measure we might have in mind. Vectors with the same number of total counts terminate along a diagonal (Fig. 2), not along a circle, meaning their Euclidean distances from the origin, could they be calculated, would be non-equal. Instead of distance from the origin, we have another straightforward measure of vector magnitude: the total number of counts, which is the number of steps from the origin to the vector terminus. This is equivalent to the Manhattan distance from the origin to the terminus.

### 3.2 Operations in count-space

The restricted algebra of semirings gives rise to operations that have clear, intuitive, physical interpretations for count data. Addition of two cell vectors produces a new vector where each element is the sum of the corresponding elements in the original vectors. This operation is the equivalent to the physical act of pooling transcripts from two cells. Adding all cell vectors is equivalent to pooling all cells into a single bulk-tissue RNA-seq experiment.

Multiplying two cell vectors also has a clear physical interpretation. The inner product, also known as the dot product, is an assessment of the similarity of two vectors based on the distribution of counts, and can be used to compare two cells. The dot product of two vectors is calculated by multiplying each corresponding element between the vectors and summing the resulting products. Because the dot product is calculated using only the operations of addition and multiplication, the natural numbers are closed under this operation.

The dot product is a measure of cell-cell transcriptome similarity, scaled by sequencing depth. A dot product of zero occurs when two cell vectors have no counts in common for any gene (the cell vectors are orthogonal). If two vectors have a large dot product, this indicates that they are both long and have many counts in common. Small dot products indicate either long vectors that have few counts in common, or short vectors for which similarity cannot be reliably determined.

Using the dot product as an indicator of cell-cell transcriptome similarity obviates many of the rationales for normalizing and transforming count data. A driving motivation behind these transformation steps is to compare distances between cells while accounting for sequencing depth. The dot product instead scales with sequencing depth, using more counts as a measure of confidence in the similarity of two cells. In other words, with more counts, we can better assess whether the RNA composition of a cell is similar or divergent from any other. As the total read count for a cell increases, the dot product increases linearly. Because of this, the dot product can be interpreted as a relative measure of similarity. Dot products can be used to find the most and least similar pairs of cells, but to interpret the absolute value of the dot product in context requires the original total number of counts.

### 3.3 Subsampling and subspaces

It is not always necessary to standardize sequencing depth across cells in order to make useful comparisons across cells, as we demonstrate in our results below. However there are certain comparisons where we might expect heterogeneity in sequencing depth to obscure biological differences, for example, when calculating differential gene expression across cells^12^. One method of standardizing sequencing depth that is coherent with our physical understanding of the measurement process is to subsample counts to an equivalent number of counts for all cell vectors. Because the scRNA-seq counting process stops at an arbitrary point, we can randomly subsample from our observations per cell to stop the process at a specific number of our choosing. By subsampling counts to the same number across cells, we can standardize sequencing depth without converting counts to proportions.

There are also cases where we might expect heterogeneity in the magnitude of expression across genes to obscure biological variation. The dot product is calculated by summing the product of each gene, meaning genes with substantially larger count values will contribute more to the dot product than genes with smaller values. This feature of the dot product can be useful for emphasizing genes with the largest dynamic range of counts, given that expression variance scales with magnitude^12^. But when highly expressed genes do not contain informative biological signal, this variance might drown out signal from genes with lower expression magnitudes. Accounting for the mean-variance relationship is the driving motivation behind several common transformations in scRNA-seq workflows. A count-based approach for tuning down the signal of highly expressed genes is to limit their total counts by randomly subsampling observations per gene. We can accomplish this by establishing a threshold for total counts and subsampling that number of observations from any gene vector that exceeds that threshold.

An alternative approach is to focus on only subsets of genes or cells when making comparisons. Subsetting the count matrix is equivalent to projecting the data onto a subspace of the original count-space. This can be conceptualized as multiplying count values for certain cells or genes by zero (or alternatively, subsampling them to zero), and is permitted using our restricted algebra. Projections onto one feature axis allow us to consider counts in only one gene, and are useful when dealing with questions about gene-level differences, such as finding genes with differential or highly variable expression. Projections onto vectors, such as projecting one cell vector onto another, are not generally possible because such a projection could result in non-natural coordinates (see Appendix 1).

With both subsampling or subsetting, there is a trade-off between reducing unwanted variation and throwing away data. Discarding counts can seem particularly painful given the already sparse nature of scRNA-seq data. We note that the standard scRNA-seq workflow already invokes subsetting by using only the top fraction of genes, ranked by variance of transformed counts. In our recommended count-based workflow, subsampling and subsetting are optional steps that should be applied depending on the research question. Furthermore, since subsampling is stochastically accomplished, we can leverage the original data by repeatedly sampling and reanalyzing to test whether results are robust to sampling effects.

## 4 countland: a count-based approach to RNA analysis

This restricted algebra can be used to achieve all the standard analytical objectives of scRNA-seq without invoking normalization or transformation (Fig. 3). We have implemented these in our software package called `countland,` written for both `python` and `R.`

**Figure 3:**
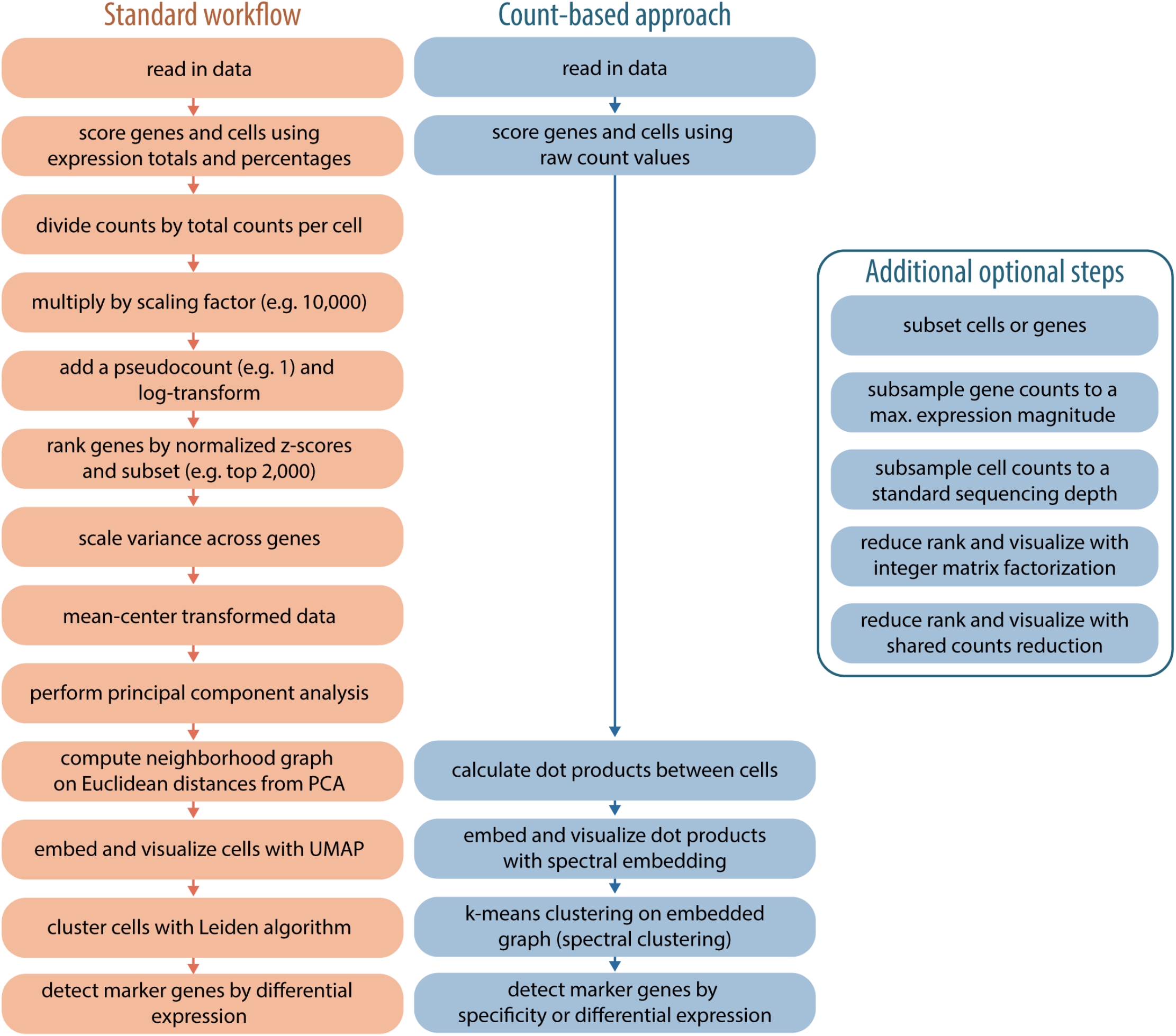
Standard and count-based workflows. A standard scRNA-seq workflow, as described in the documentation for `scanpy` and `Seurat` is shown in orange. A count-based approach, as implemented in `countland,` is shown in blue, with optional subsetting, subsampling, and rank reduction steps adjacent.

### 4.1 Comparing cells

A common approach for comparing cells in standard, normalization-based scRNA-seq workflows is to use principal component analysis (PCA) to rotate and project expression values onto a smaller dimensional space, and then calculate a neighborhood graph using the cell-cell distances in the PCA representation^6^. In a count-based approach, we avoid the use of distances by calculating the dot product of untransformed counts between all pairwise combinations of cells. This process generates a similarity matrix, rather than a distance matrix. This similarity matrix has several important properties: it is square, with the number of rows and columns determined by the number of cells, and it is symmetric, with diagonal values giving the dot product of each vector with itself, which is the sum of squares of all its elements. Off-diagonal values are the dot products between different cell vectors, and these values range from zero to any positive integer, unbounded on the maximum end as alignment increases.

Cell similarity can be visualized by applying spectral embedding to this matrix. Spectral embedding involves calculating the graph Laplacian of the matrix and then estimating the eigenvectors and eigenvalues of this graph^21^. The cell similarity matrix can be visualized in two dimensions by plotting points using two eigenvectors (Fig. 4). This matrix can also be used to cluster cells by applying a k-means algorithm to the eigenvector matrix (spectral clustering). Here we demonstrate the use of spectral clustering of the dot product matrix for identifying cell populations using the Gold standard scRNA-seq benchmark dataset, described by Freytag *et al* (2018)^16^. This dataset consists of 925 cells from three populations derived from human lung tissue. Using `countland,` we recover these populations with high fidelity (adjusted rand index = 0.994, see benchmark results below). While spectral embedding includes operations that are not closed for natural numbers (e.g. subtraction and division), here they are operating on comparisons between cells (the dot product matrix) that do respect the count nature of the data.

**Figure 4:**
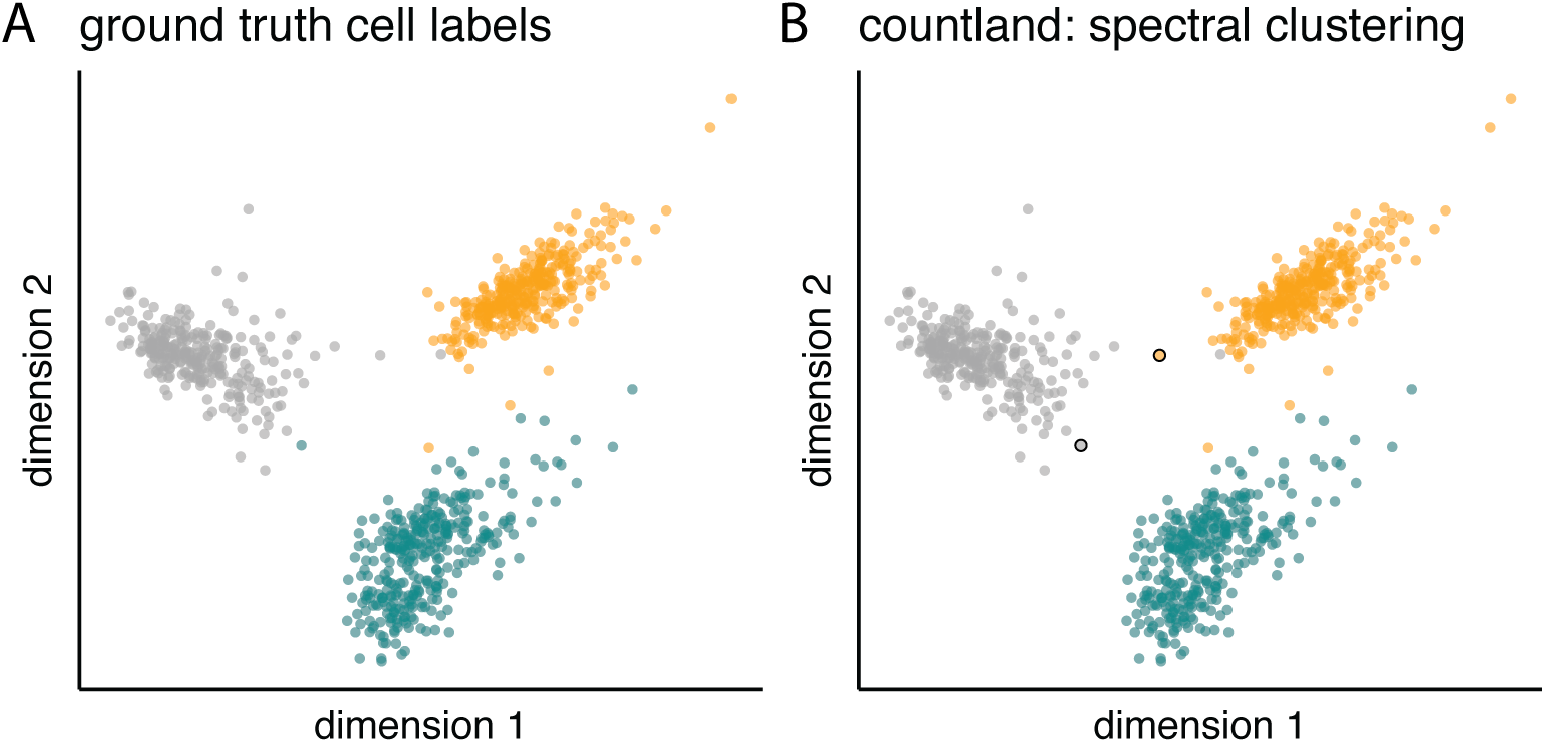
Clustering cells by similarity using a count-based approach. The Gold standard scRNA-seq dataset of 925 cells from human lungs are visualized using spectral embedding of the pairwise dot product matrix between all cells. A, cells are colored according to the ground truth labels as described in Freytag *et al* (2018)^16^. B, cells are colored according to the results of spectral clustering. The two points outlined in black are the only cells for which cluster labels differed from ground-truth labels.

### 4.2 Comparing genes

Gene expression can be compared with a count-based approach as well. There are several count-based measures that provide insight into expression dynamics. These include:

- The total number of counts per gene, summed across cells.
- The maximum observed count value per gene.
- The number of cells where a gene was detected, as well as the number where detected at a count value larger than one or some other value.
- The number of unique count values (e.g. 0, 1, 2). Given the discrete nature of low-magnitude count values, this measure can provide insight into expression variability across cells.
- The largest number *n*, where there are *n* cells with ≥*n* counts for the gene in question. We refer to this measure as the *count index*, and it can be helpful for finding genes that frequently show higher count values, as compared to genes that are mostly detected at values of 1 or 2, with a few high-count exceptions.

The ideal marker gene for a cluster of cells can be defined as the gene with the highest differential expression between cells in the cluster and all other cells, or in an alternative approach, as the gene that is most specifically expressed in cluster cells^12^. A count-based approach for identifying marker genes by specificity is to count the number of cells with non-zero observations for each gene, and then calculating the difference between the fraction of these cells in a cluster versus the fraction that are not. The ideal marker gene would return a value of one, indicating it was expressed in all cluster cells and no others.

Differential expression can be assessed using rank-sums tests, similar to those invoked in standard workflows, but here applied to raw counts instead of transformed proportions. In this method, counts for a given gene are ranked between cluster and non-cluster cells, the ranks for each group are summed, and a test statistic is calculated. This statistic is used to test the hypothesis that observations from cluster cells are larger than those from non-cluster cells. Because heterogeneity in count depth can influence the magnitude of expression for individual gene, subsampling to a standard sequencing depth prior to calculating differential gene expression is recommended.

### 4.3 Reducing dimensionality and low-rank approximation

The high dimensionality of single-cell count matrices is one of the fundamental challenges to analyzing, interpreting, and visualizing these data^22^. There are several motives for embedding the count matrix in a lower-dimensional space, primary among them being data visualization. In the standard workflow, dimensional reduction is achieved through linear transformation (e.g. PCA and projection) of the already-transformed count matrix, followed by further non-linear reductions to two-dimensions using t-SNE or UMAP (Fig. 3). Recent reports suggest that rather than reducing the distances between similar cells, this approach results in extensive distortions of cell-cell distances^22^.

Here we implement two count-based methods for reducing the number of dimensions that are not based on Euclidean distance. One way of reducing the dimensionality of count data is to combine genes with similar information content. We measure this as the number of shared counts between two genes; this is calculated by comparing two gene vectors (e.g. matrix rows), taking the smaller of the two count values for each cell (e.g. each column), and summing. This sum represents the number of times both genes were counted in the same cell. Calculating this sum for all pairwise combinations of genes results in a similarity matrix of shared counts. To find cohorts of genes that have relatively large numbers of shared counts, we perform spectral clustering on this matrix. We use these clusters to reduce the dimensionality of the count matrix by pooling counts (summing) across genes in the same clusters. This method reduces a matrix with the dimensions *m* cells by *n* genes to one with the dimensions *m* cells and *k* meta-genes, where *k* depends on the number of clusters found using spectral clustering. Cells can be visualized using this approach by plotting their coordinates along a pair of meta-genes (Fig. 5B).

**Figure 5:**
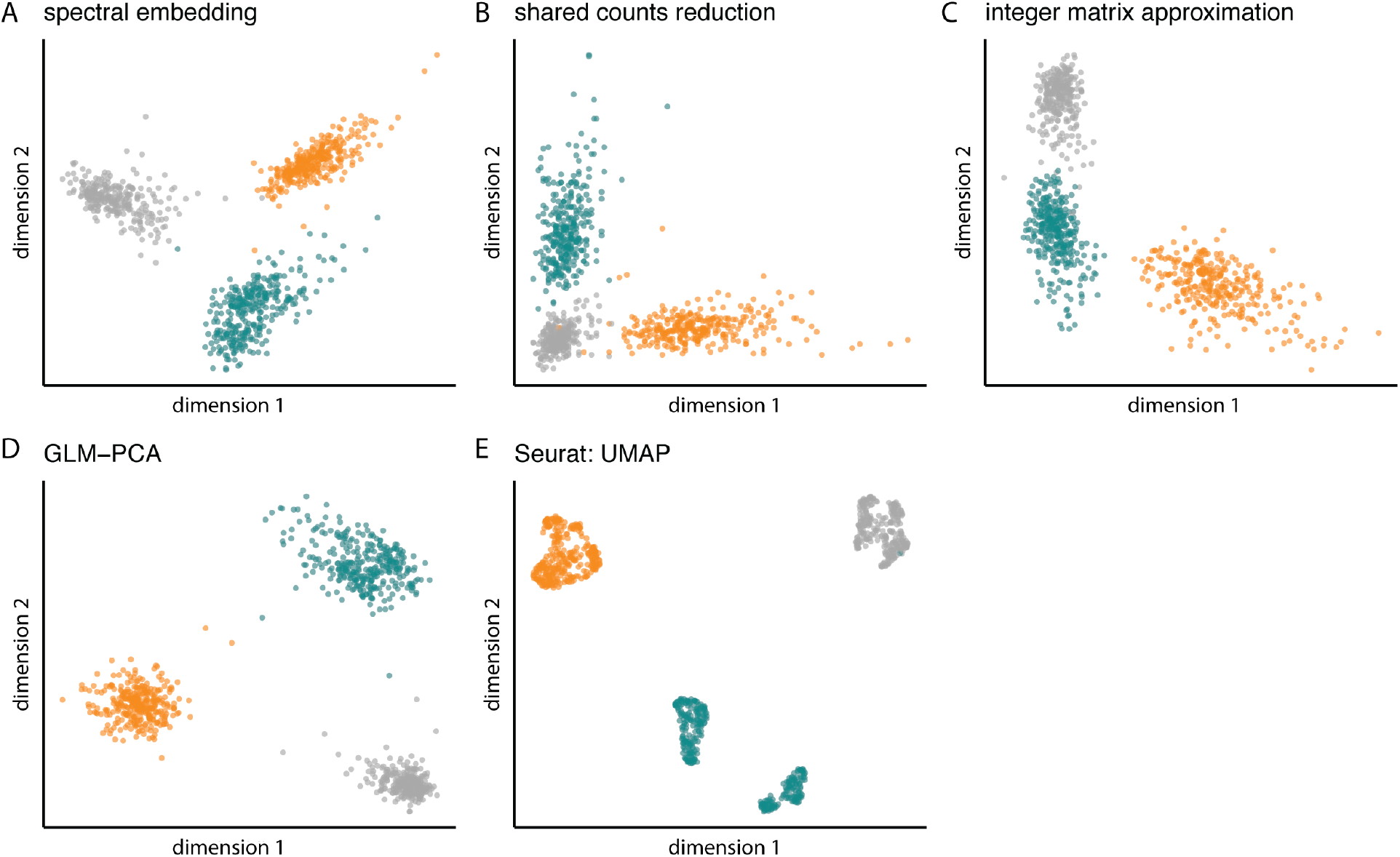
Comparing dimensional reduction approaches. Cells from the Gold standard benchmark dataset, colored in all panels according to ground truth labels. A, spectral embedding of the dot product matrix. B, reduced dimensionality by pooling genes with large numbers of shared counts. C, reduced dimensionality using integer matrix approximation. D, generalized PCA (GLM-PCA), as implemented in the R package `glmpca`. ^7^ E, uniform manifold approximation (UMAP) following data transformation, as implemented in `Seurat.` Note that UMAP on transformed values results in the most apparent cluster boundaries, but it also results in one ground truth population (cyan) being split into two groups (see performance evaluation below).

Integer matrix factorization is an alternative approach to achieve a low-rank approximation of matrices that include only natural numbers^23^. Like other matrix factorizations (e.g. singular-value decomposition), this method seeks to find lower-rank matrices that can be multiplied together to approximate a higher-rank matrix, here the count matrix. Integer matrix approximation generates three matrices, termed **U, V**, and **Λ**, similar to the three matrices generated by singular-value decomposition on data consisting of real numbers. When using integer matrix approximation on single-cell count data, matrix **U** has the dimensions m cells by *k* features, with *k* provided as the target rank, **V** has the dimensions *k* features by *n* genes, and **Λ** is a diagonal matrix of *k* scaling factors. Because of the discrete nature of count data, this factorization cannot be accomplished conventionally, but approximations for this factorization have been proposed for other types of count-based data^24^. Here we implement the algorithm for integer matrix approximation in python and R, and apply it to the approximation of single-cell count data. Following integer matrix approximation, cells can be embedded in a lower-dimensional space by multiplying the count matrix with matrix **V**, scaled by **Λ**. Cells can then be visualized by plotting their coordinates in two resulting embedding components (Fig. 5C). This is conceptually similar to calculating principal components for visualization using singular-value decomposition.

Given the problems inherent to data transformation, Townes *et al* (2019)^7^ recently proposed a generalized version of PCA (GLM-PCA) that takes advantage of the exponential family of likelihoods, and which can be applied directly to raw counts. This method, like those described above, also does not rely on Euclidean distances, but unlike the shared counts and integer matrix approximation approaches, it does not preserve the count-like nature of the data (values will not necessarily be natural numbers). Given that it allows for visualization without transformation, however, we highlight it as a promising additional avenue for interpreting scRNA-seq data.

## 5 Evaluating count-based approaches to clustering

### 5.1 Gold standard

We tested the `countland` methods described above on published benchmark data to evaluate its performance relative to ground truth and consistency with published standard workflows. As shown in Figure 3, `countland` recovers the ground truth cell identities from the Gold standard dataset with high fidelity. Clustering accuracy was evaluated using three measures: the adjusted rand index (ARI), normalized mutual information (NMI), and cluster homogeneity. ARI and NMI both evaluate the similarity between two sets of clusterings, while homogeneity measures the degree to which an identified cluster contains only members of one ground truth group. Against ground truth, `countland` returns an ARI score of 0.994, an NMI of 0.987, and homogeneity of 0.987. These results were achieved using `countland` on the raw count matrix, demonstrating that without any form of normalization or data transformation, count-based methods can accurately identify cells by expression.

For the same dataset, `Seurat` returns a lower ARI (0.409) when using the default parameters, but after reducing the resolution used in clustering, `Seurat` returns an ARI of 0.997. The difference in scores with `Seurat` can be attributed to the software splitting one ground truth cell population into two clusters, as seen in the cyan colored cells in Fig. 5E.

Because the Gold standard dataset is significantly less sparse than many scRNA-seq datasets, we tested `countland’s` performance on a modified Gold standard dataset where each cell contained only 1% of the original number of observations. Both `countland` and `Seurat` can recover ground truth cell identities with high fidelity (ARI 0.997 and 0.997, NMI 0.993 and 0.993, homogeneity 0.993 and 0.994, respectively). We note that when analyzing the more sparse dataset, `countland` achieves slightly better results when the number of components (Laplacian eigenvectors) used in spectral clustering are increased from 5 to 10 (ARI increased from 0.977 to 0.99).

We tested performance on datasets derived from the Gold standard, modified to introduce the kinds of data heterogeneity that are often invoked to justify data transformations. First, we modified the sparse Gold standard dataset so that 100 of the 925 cells had their original measurements from before reducing counts to 1% of their original number. These 100 cells were randomly drawn 50 each from two of the three cell populations. With this dataset we observed a reduction in cluster accuracy analyzing counts directly (ARI 0.399, NMI 0.431, homogeneity 0.431). But after applying `countland’s` subsampling procedure to bring cells to a standardized sequencing depth, `countland` accurately recovers ground truth cell identities (ARI 0.952, NMI 0.927, homogeneity 0.927). In contrast, `Seurat` fails to accurately identify cells, despite normalizing data by sequencing depth (ARI 0.478, NMI 0.555, homogeneity 0.555). Visualizing the `Seurat` results shows that clusters separate cells by sequencing depth as well as their original identities. This is likely because cells with more total counts have observations for many genes that are not observed in lower-count cells, a fact which depth normalization cannot account for.

We also tested performance on a version of the sparse Gold standard dataset modified to have substantial heterogeneity in gene expression. To accomplish this we added ten highly expressed genes with no variation attributed to cell population. Count values were simulated for these genes using a Poisson distribution with a lambda value 10x larger than the largest observed mean count value across genes. Adding these genes with high count values resulted in decreased accuracy for `countland` when analyzing raw counts (ARI 0.347, NMI 0.408, homogeneity 0.408), but this effect was eliminated when highly expressed genes were subsampled to a predetermined maximum count value (ARI 0.993, NMI 0.987, homogeneity 0.987). The performance of `Seurat` was robust to the addition of highly expressed genes (ARI 0.997, NMI 0.993, homogeneity 0.993).

### 5.2 Silver standard

We evaluated the performance of `countland` and `Seurat` on the Silver standard dataset (version 3a) of peripheral mononuclear cells (PBMCs). This dataset does not include ground truth labels, instead it includes cell labels derived from similarity to a reference dataset^25^. Previous evaluations of scRNA-seq analysis software show that all methods fail to recover all labeled cell groups, with different methods returning an ARI between ∼0.2 and ∼0.6. We reanalyzed this dataset using `Seurat` and observed an ARI of 0.456, NMI of 0.622, and homogeneity of 0.679.

When subsampling highly expressed genes to a maximum total count value equal to the number of cells, countland returned similar results to other clustering software (ARI 0.442, NMI 0.594, homogeneity 0.594). Subsampling genes even further returned higher scores (ARI 0.534, NMI 0.633, homogeneity 0.633), as did subsetting the data to remove the top 5% of genes altogether (ARI 0.57, NMI 0.644, homogeneity 0.644). This suggests that, for the Silver standard dataset, highly expressed genes are not useful in identifying cells according to the published labels, and may mask signal present in other genes. Subsampling cells to a standard sequencing depth did not result in an increase in clustering scores (ARI 0.34, NMI 0.576, homogeneity 0.576).

## 6 Discussion

The best way forward can sometimes require first taking a few steps back. Measurement theory^5^ provides a roadmap for this in the context of scRNA-seq data analysis: start by considering the measurement process by which numbers are assigned to attributes, and then consider how mathematical operations on the measurements correspond to physical processes. This is fundamental to understanding how our measurements address our research question. This approach is especially valuable for navigating the data-rich world of next generation sequencing and functional genomics. Current practices using these high-dimensional data almost always involve *ad hoc* transformations that are applied with a general sense that something must be done to the data before downstream analyses are possible (e.g. the data must be transformed so that Euclidean distances become meaningful). We advocate for a more intentional approach, where data processing steps are only taken when they respect the measurement process and will be informative for the research question.

Current practices in single cell analysis have been highly optimized with one primary objective: to identify and visualize clusters of cells. But modern research with scRNA-seq data has far more ambitious and diverse objectives, including identifying developmental trajectories, comparing cells and genes across species and evolutionary time, and associating expression differences with phenotypes of interest. The result is that a substantial research effort is currently dedicated to things other than clustering, and this requires undoing many problems created by data preprocessing steps that were developed with clustering in mind. Here we have described an approach that seeks to sidestep, rather than patch, many current practices in order to avoid these problems altogether.

Our results show that the restricted algebra of count-space is sufficient to perform many common scRNA-seq analysis tasks, while respecting the underlying count nature of the data. For example, assessing cell similarity with the dot product of transcript counts is a powerful tool to identify distinct clusters that correspond to real cell populations (Fig. 3). The restricted algebra implemented in `countland` works well, even when datasets are sparse, without taking any steps to account for heterogeneity in sequencing depth or gene expression magnitude. This indicates that it is not universally necessary to convert counts to fractional abundances or log-transform values in order to categorize cells by expression. And in cases where heterogeneity in those measures obscures biological differences, we provide a count-based solution via subsampling, and demonstrate that this solution can match or outperform standard approaches.

However, we anticipate that there are scenarios when the standard, transformation based approach can recover clusters that are not identified using count-based approaches. There are almost certainly circumstances in which these transformations have the effect of exaggerating subtle but perhaps real biological differences between cell populations. The challenge is that these circumstances would be indistinguishable from situations where the same transformations instead result in exaggerating spurious and artifactual population structure. Count-based approaches, on the other hand, are easy to interpret; more similar cells are those that have more transcripts in common.

There is great potential for improvements in count-based approaches, especially in the area of spectral embedding and clustering of the dot product matrix. While standard, distance-based clustering methods have undergone many generations of improvements, here we have used an out-of-the-box approach to spectral clustering, as implemented in `scikit-learn`, a standard clustering library in `python`^26^ (which we reimplemented in `R`). Future optimizations to the choice of graph Laplacian or the clustering algorithm that are specific to single-cell data may result in improved performance in identifying cell populations using transcript counts. However, we stress that segregating and labeling cells is only one small portion of the analyses that are possible with scRNA-seq data. Cells do not always fall into clear populations that will conform to clustering algorithms, especially when considering cells over their developmental lifetimes. The dot product is an intuitive and powerful measure of cell-cell similarity, even when not invoking clustering. Comparisons of cells across timepoints, treatments, tissues, and species are all facilitated by leveraging a count-based approach that avoids *ad hoc* transformations and improves interpretability.

## 7 Acknowledgments

We thank Abby Skwara, Kevin O’Neil, Milo S. Johnson, Seth Donoughe, and members of the Dunn lab for comments on early drafts of this manuscript. This material is based upon work supported by the NSF Postdoctoral Research Fellowship in Biology under Grant No. 2109502.

## 8 Methods

### 8.1 countland

We implemented `countland` in `python` and `R,` available at https://github.com/shchurch/countland. Implementation in both languages will allow these methods to be more widely used, and will provide an opportunity to cross validate results.

The implementation of the restricted algebra in our package `countland` is summarized here using mathematical notation. See Appendix 1 for a background on the mathematical principles that justify our restricted algebra, and the git repository for more specifics.

Let **C** be the count matrix containing the raw measurements, and let N_0_ be the set of natural numbers, inclusive of 0, where **C**_*ij*_ ∈N_0_. **C** has the dimensions *m* cells and *n* genes. The established convention for **C** in python has cells as matrix rows and genes as columns, whereas in R, genes are rows and cells are columns. Our descriptions in the text of this manuscript use the `python` convention, unless stated otherwise.

#### 8.1.1 Dot product matrix

Let **D** be the *m* × *m* dot product matrix, where element **D**_*ij*_ is the dot product of cell *i* with cell *j*. **D** is calculated using matrix multiplication as **D** = **CC**^*T*^. Because this requires only multiplication and addition operations on the elements, this does not move the data out of count-space and will return only values contained in N_0_. Note that this is conceptually very similar to the calculation of the covariance matrix, which is at the heart of methods often used in scRNA-seq analyses, such as PCA. Covariance matrices are calculated as **XX**^*T*^, where **X** is the mean-centered transformation of **C**. However calculating the mean and centering requires division and subtraction and moves the data out of count-space, resulting in negative and fractional values not contained in N_0_.

#### 8.1.2 Spectral clustering

Spectral embedding and clustering require a similarity matrix as input. Here, **D** is used as the cell-cell similarity matrix. Prior to spectral embedding, the diagonal elements of **D** are replaced with zeros to remove edges of the similarity graph that connect cells to themselves.

In the python implementation of `countland,` we used spectral clustering functions as written in scikit learn, modified so that it returns the eigenvalues as well as the eigenvectors of the graph Laplacian. As with `scikit learn,` in `countland` the user can input the target number of clusters as well as the number of components (eigenvectors of the graph Laplacian) that should be considered in the spectral clustering algorithm.

To ensure that methods and results are directly comparable between the `python` and `R` versions of countland, instead of using an existing package for spectral clustering (e.g. R:Spectrum^27^), we re-implemented the `scikit learn` algorithm from scratch in `R.`

To facilitate the choice of the number of clusters, we implemented a common heuristic for determining the optimal numbers of clusters by considering the difference between eigenvalues *λ*_1_, …, *λ*_*n*_ of the graph Laplacian. According to this heuristic, a reasonable choice for the optimal number of clusters *k* is one where the eigenvalues *λ*_1_, …, *λ*_*k*_ are relatively small, and the gap |*λ*_*k*+1_ −*λ*_*k*_| is relatively large. This heuristic is intended as a guideline, and other methods of selecting *k* may also be useful (e.g. using *a priori* biological information).

#### 8.1.3 Subsampling counts

Sequencing depth in scRNA-seq experiments is not standardized across samples, with the result that the sum of counts per cell 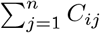 varies. Sequencing depth can be standardized in `countland` by subsampling observations to a fixed number *x* of total counts per cell, such that 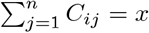. This is accomplished by flattening each cell vector to an array of transcripts, with the frequency of each transcript given by its count value, and then randomly choosing *x* transcripts without replacement. The new cell vector is given by the sampled transcript frequency.

A similar approach is taken when subsampling genes to reduce the impact of heterogeneity in expression magnitude. In this case, each gene vector is flattened to an array of cell observations. For any gene with *> x* total observations, *x* observations are randomly chosen without replacement, and the new gene vector is given by the sampled cell frequency.

#### 8.1.4 Expression scores

The following expression scores for cell *i* can be calculated in `countland:`

- total counts 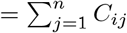
- maximum count value = max_1≤*j*≤*n*_(*C*_*ij*_)
- The number of observations above zero 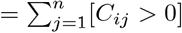 where [*F*] = 0 if *F* is false, and 1 if *F* is true
- The number of observations above one 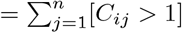
- The number of observations above ten 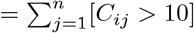
- The number of unique count values = unique_1≤*j*≤*n*_(*C*_*ij*_)
- The count index *c* = max_1≤*j*≤*n*_(*C*_*ij*_) where 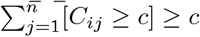

As well as the corresponding scores along the *n* dimension.

We implemented two methods for comparing differential expression across clusters. Let *M* ^′^ be the cells within the cluster and *M*^′′^ be the rest. For the first method, we calculate the difference between proportions of non-zero observations

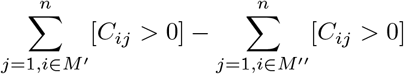

For the second method, we use a Wilcoxon Rank-sum test (Mann Whitney U test) between *M* ^′^ and *M* ^′′^, followed by Benjamini-Hochberg false discovery rate correction of p-values (*α* for significance set at 0.05).

#### 8.1.5 Dimensional reduction via shared counts

Let **S** be the *n* × *n* matrix of shared counts between genes, where element **S**_*pq*_ is calculated as 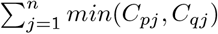 *min* (*C*_*pj*_, *C*_*qj*_). **S**_*pq*_ is equal to 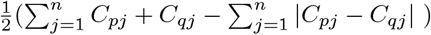 where 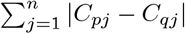 is the Manhattan distance.

**S** is a similarity matrix; it describes the amount of shared information between pairs of genes. In `countland` we apply spectral clustering to **S** to identify clusters of genes with strong signatures of shared information. The number of clusters *k* is determined by the user and the number of components considered is set equal to the number of clusters. Let *N* ^′^ be the genes in an identified cluster and let **C**^′^ be the reduced *k* × *m* count matrix, the values of the reduced count matrix 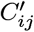 are calculated as 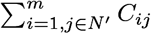

#### 8.1.6 Integer matrix approximation

Integer matrix factorization is a method for estimating lower rank matrices that can be multiplied to approximate a higher rank matrix that contains discrete values, such as integers^23^. This factorization generates three matrices, **U, V**, and **Λ. U** has the dimensions *m* × *k* features, with *k* provided by the user. **V** has the dimensions *k* × *n* genes, and **Λ** is a *k* × *k* diagonal matrix of scaling factors. This is conceptually similar to other matrix factorizations (e.g. singular-value decomposition) that generated matrices **U, V**, and **Σ**.

An approximation for integer matrix factorization has been implemented for `MATLAB` in the application `SUSTain`^24^. We re-implemented the required functions for integer matrix approximation on a matrix in `python` and `R,` and have made them available for public use at https://github.com/shchurch/integer_matrix_approximation.

As in `SUSTain,` integer matrix approximation is accomplished in three steps. First, parameters are set, including: the target rank of the factorized matrices, the upper and lower bounds of the integer values (default lower bound is zero), the maximum number of iterations (default is 1000000), and the stopping criterion (default is a difference of 0.0001). Second, initial matrices **U, V**, and **Λ** are calculated by sampling integers from the higher rank matrix, ensuring that values remain within the bounds. Third, **U, V**, and **Λ** are updated via the algorithm described in the corresponding `SUSTain` manuscript^24^.

### 8.2 Performance evaluation

We evaluated `countland’s` performance using the Gold and Silver standard benchmark scRNA-seq datasets provided by Freytag *et al* (2018)^16^. The Gold standard dataset includes data from 925 cells drawn from three populations of human lung adenocarcinoma lines. This dataset has ground truth cell labels, but is far less sparse than most scRNA-seq datasets (29% of values are non-zero). To evaluate performance on a more sparse dataset, we created a modified Gold standard dataset where each cell contained only 1% of its original number of observations.

For the Gold standard dataset, we evaluated performance after increasing the amount of heterogeneity in sequencing depth and in gene expression. To manipulate heterogeneity in sequencing depth, we restored 100 cells from the sparse Gold standard dataset to their original sequencing depth, drawing 50 cells each from two of the three cell populations present in the data. To manipulate heterogeneity in gene expression, we simulated counts for 10 genes, using a Poisson distribution with lambda values 10x larger than the largest observed mean count value across genes.

The Silver standard datasets are composed of fresh peripheral mononuclear cells (PBMCs). Here we have evaluated performance on dataset 3a, named by the original authors, which contains 4,310 cells labeled by matching cells to a reference dataset by expression. These labels do not constitute a ground truth but have been shown to match identities found using marker genes to classify cells^25^.

For the Gold and Silver standard datasets, we tested `countland` first on raw count matrices, meaning no subsampling or subsetting was performed prior to calculating the dot product and performing spectral embedding and clustering. We then tested the impact of subsampling cells to a standard sequencing depth, and of subsampling genes to a maximum total count value. In the case of the Silver standard dataset, we tested two thresholds for maximum gene counts: a threshold equal to the number of cells, and one equal to 1/2 the number of cells. We also tested the effects of dropping the top 5% of genes according to total counts. For the Gold standard dataset, we tested the impact of varying the number of components (Laplacian eigenvectors) used to identify clusters.

All datasets and code required to perform analyses are provided along with the software at github.com/shchurch/countland.

### 8.3 Data availability

The countland package for `python` and `R`, as well as all data and code required to reproduce the results shown here is available at https://github.com/shchurch/countland.

## 9 Appendix 1. Properties of vector spaces over zero and the natural numbers

This appendix provides additional background and information on mathematical concepts relevant to the analysis of transcript count matrices.

### 9.1 The natural numbers, N_0_

The natural numbers are a group of all positive integers. Depending on the definition, this group may also include zero (i.e. all non-negative integers), a convention we follow here so that N_0_={0,1,2,3,… }. We can classify groups of numbers by the operations under which they are closed, meaning operations on elements in the group result in elements that are also in the group. The most familiar group is a field, which is closed under addition, subtraction, multiplication, and addition, and includes rational numbers Q and real numbers R. The natural numbers form a semiring, because N_0_ is closed for multiplication and addition, but not subtraction or division.

N_0_ contains the additive identity element, 0, that can be added to any element to return the same element (e.g. *x* + 0 = *x*). N_0_ also contains the multiplicative identity element, 1, that can be multiplied with any element to return the same element (e.g. *x*1 = *x*).

Inverses are elements that return the identity elements under specified operations. For example, the negative numbers are inverses under addition, because *x* + (−*x*) returns the additive identity element, 0. However, because N_0_ does not contain negative numbers, it doesn’t have additive inverses. Similarly, reciprocal values are inverses under multiplication, because *x*(1*/x*) returns the multiplicative identity element, 1. N_0_ likewise does not contain reciprocals, so therefore doesn’t have multiplicative inverses.

Subtraction can be defined as the addition of an additive inverse (a negative number), and division can be defined as multiplication with a multiplicative inverse (a reciprocal). Because these two inverses are not contained in N_0_, there is no subtraction or division. This is equivalent to the observation that N_0_ is not closed for subtraction or division.

### 9.2 Vector spaces over N_0_

We can build the vector space over N_0_ as the group of all vectors **V** such that

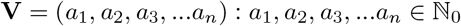

The vector space over N_0_ can be envisioned as being restricted to the integer grid that is located over the upper right quadrant of a coordinate system, inclusive of the origin and axes. Certain operations are possible in this restricted space, while others are not. For example, we can apply the operation of the inner product (dot product) because this operation requires only multiplication and addition of vector elements. However, unlike a vector space over a field, in the vector space over N_0_ there are no angles between vectors. Calculating angles from dot product requires division by vector length, and N_0_ is not closed for division.

Furthermore, in count-space, vector length is not a Euclidean measure of distance as there is no equivalent measure of distance in a space without subtraction or square roots. Instead of Euclidean distance, we can use the number of integer steps in a positive direction as a measure of length, which is equivalent to the Manhattan distance between the vector terminus and the origin.

Vector rotation is not possible in count-space as it would require rotation matrices with new basis vectors that include negative elements. Vector reflections are possible, because we are free to permute our count matrix, as are shears of vectors. Some vector projections are possible, but not all. For example, if we project vector **b** onto vector **a**, the result will be a multiple *q* of **a**, *q***a**. The value of that multiple will be equal to *q* = (**a**^*T*^ **b**)*/*(**a**^*T*^ **a**), which requires division to calculate unless **a**^*T*^ **a** = 1. Over N_0_, that only happens when there is a single entry that is 1, i.e. when vector **a** is one of the original basis vectors. Therefore we can project vectors onto basis vectors, but not onto arbitrary vectors (e.g. we cannot project one cell vector onto another). Projecting onto basis vectors is the equivalent of multiplying some values by 0 while retaining others.

Without rotation and vector projection, it is clear that certain complex operations like principal component analysis that rely on these are not possible in this vector space.

### 9.3 High-dimensional, low-magnitude vector spaces over N_0_

While the above pertains to vector spaces over N_0_ in general, there are interesting properties of the very high dimensional, low magnitude vector spaces that describe scRNA-seq count data.

Most gene expression datasets contain measurements for many thousands of genes, meaning this vector space has many thousands of dimensions. Furthermore, due to the sparse nature of these count matrices, it is difficult if not impossible to find a reasonable lower-dimensional approximation. In other words, because many features contain only a few, non-overlapping observations, there is no way to reduce the rank of this matrix without discarding features.

Because most measures of gene expression are low-magnitude integers, most cell vectors terminate only a few steps from the origin in any given direction. This does not mean that vectors are close to the origin overall. Vector length is non-Euclidean; it is calculated as the sum of steps back to the origin (Manhattan distance), not the distance along a diagonal (Euclidean distance).

Because the vast majority of values in the count matrix are zero, cell vectors are perpendicular to each other in many directions. The may result in the outcome that cell-cell similarity has more to do with the number and distribution of non-zero observations than expression magnitude. This has been demonstrated by the fact that binary transformations of scRNA-seq data to zero/non-zero contain enough information to recapitulate major patterns in the data^11^.

